# Genome-wide patterns of non-coding sequence variation in the major fungal pathogen *Aspergillus fumigatus*

**DOI:** 10.1101/2024.01.08.574724

**Authors:** Alec Brown, Jacob L. Steenwyk, Antonis Rokas

**Author notes:** Address correspondence to: Antonis Rokas.

## Abstract

*A. fumigatus* is a deadly fungal pathogen, responsible for >400,000 infections/year and high mortality rates. *A. fumigatus* strains exhibit variation in infection-relevant traits, including in their virulence. However, most *A. fumigatus* protein-coding genes, including those that modulate its virulence, are shared between *A. fumigatus* strains and closely related non-pathogenic relatives. We hypothesized that *A. fumigatus* genes exhibit substantial genetic variation in the non-coding regions immediately upstream to the start codons of genes, which could reflect differences in gene regulation between strains. To begin testing this hypothesis, we identified 5,812 single-copy orthologs across the genomes of 263 *A. fumigatus* strains. *A. fumigatus* non-coding regions showed higher levels of sequence variation compared to their corresponding protein-coding regions. Specifically, we found that 1,274 non-coding regions exhibited <75% nucleotide sequence similarity (compared to 928 protein-coding regions) and 3,721 non-coding regions exhibited between 75% and 99% similarity (compared to 2,482 protein-coding regions) across strains. Only 817 non-coding regions exhibited ≥99% sequence similarity compared to 2,402 protein-coding regions. By examining 2,482 genes whose protein-coding sequence identity scores ranged between 75% and 99%, we identified 478 total genes with signatures of positive selection only in their non-coding regions and 65 total genes with signatures only in their protein-coding regions. 28 of the 478 non-coding regions and 5 of the 65 protein-coding regions under selection are associated with genes known to modulate *A. fumigatus* virulence. Non-coding region variation between *A. fumigatus* strains included single nucleotide polymorphisms and insertions or deletions of at least a few nucleotides. These results show that non-coding regions of *A. fumigatus* genes harbor greater sequence variation than protein-coding regions, raising the hypothesis that this variation may contribute to *A. fumigatus* phenotypic heterogeneity.

## Introduction

Invasive Aspergillosis (IA) is one of the deadliest fungal diseases to humans, responsible for over 400,000 infections per year with a mortality rate of >50% (Bongomin et al., 2017). Most IA cases (>90%) are caused by *Aspergillus fumigatus* (Steinbach et al., 2012; Rokas et al., 2020), a saprophytic fungus commonly found in the soil (Flores et al., 2014) as well as urban environments, such as waste piles and hospitals (Wirmann et al., 2018). In its natural environment, *A. fumigatus* plays an important role in nitrogen and carbon recycling (Latge et al., 2019). *A. fumigatus* has adapted over time to survive environmental pressures, such as high temperatures, variation in pH, and low oxygen availability (Bhabhra & Askew, 2005; Park & Yu, 2016; Rees et al., 2017), and to compete with other microorganisms for resources (Latge et al., 2019). Recently, the World Health Organization included *A. fumigatus* in its first ever list of fungal “priority pathogens”, a testament to its seriousness as a threat to public health (WHO, 2022).

*A. fumigatus* typically reproduces via asexual spores (conidia), which are released into the air for eventual germination. While some spores eventually return to the soil, others are inhaled by humans, and interact with the epithelium of the lung (Chotirmall et al., 2013). Aided by their small diameter (2-3 µm) and hydrophobic outer layer, these spores can subsequently reach the lung alveoli (Croft et al., 2016). Once in the lung, *A. fumigatus* must survive a hostile environment and host defense system (Bertuzzi et al., 2018). Immunocompetent individuals clear these spores, but immunocompromised ones are at risk of developing IA (Cadena et al., 2021). Several species closely related to *A. fumigatus* are not considered pathogenic (de Vries et al., 2017; Rokas et al., 2020; Mead et al., 2021). For example, *Aspergillus fischeri* is a close relative of *A. fumigatus* (the two species share >90% average nucleotide sequence identity and >95% average amino acid sequence identity between orthologs), yet *A. fischeri* is less virulent and is not considered clinically relevant (Mead et al., 2019; Steenwyk et al., 2020). Early genomic comparisons between two strains of *A. fumigatus* (Af293 and A1163) and one strain of *A. fischeri* (NRRL 181) revealed a set of genes uniquely present in *A. fumigatus* (Fedorova et al., 2008). However, a more recent genomic examination of 18 *Aspergillus* section *Fumigati* strains representing 13 species found that 206 known genetic determinants of virulence in *A. fumigatus* are all shared between *A. fumigatus* and at least one other closely related, non-pathogenic species (Mead et al., 2021). Finally, recent examinations of genomic variation between the genomes of hundreds of *A. fumigatus* isolates (Barber et al., 2021; Lofgren et al., 2021, Horta et al., 2022) have revealed that *A. fumigatus* has an open pangenome with ∼70% of its genes being highly conserved across strains (core) and that orthologs from both clinical and environmental strains exhibit a high degree of sequence conservation.

Variation in non-coding regions can also contribute to phenotypic variation, including gene expression variation between and within species (Caroll, 2005; Hill et al., 2020). We have previously demonstrated that non-coding regions between two *A. fumigatus* reference strains, Af293 and A1163 (Brown et al., 2022; Colabardini et al., 2022), as well as between *A. fumigatus* and its non-pathogenic close relatives are highly variable (Brown et al., 2022). For example, we found that 418 *A. fumigatus* genes exhibit a different rate of evolution in their non-coding regions (relative to non-pathogenic close relatives), including the non-coding regions of 25 genes that are known genetic determinants of *A. fumigatus* virulence. Examination of these non-coding regions revealed numerous single nucleotide and insertion/deletion (indel) differences between *A. fumigatus* and closely related non-pathogenic species (Brown et al., 2022).

To increase our knowledge of non-coding region variation within *A. fumigatus* and how levels of non-coding sequence variation compare to levels of protein-coding sequence variation, we examined the genomes of 263 *A. fumigatus* strains (using 2 *A. fischeri* strains as an outgroup). Of the 5,812 single-copy orthologs identified across all strains, 2,402 genes had identical or near-identical sequences (≥99%); the same was true for 817 non-coding regions; 3,721 non-coding regions exhibited between 75% and 99% similarity (compared to 2,482 protein-coding regions); and 1,274 genes had a percent identity <75% in their respective non-coding regions and 928 in their protein-coding regions. Regions with low sequence similarity tend to yield unreliable sequence alignments and were not included in subsequent analyses. Instead, we focused our analyses on 2,482 genes whose protein-coding regions exhibited percent sequence identities between 75% and 99%, performing two different tests of positive selection: the McDonald-Kreitman (MK) test (McDonald & Kreitman, 1991, Murga-Moreno et al., 2019) and the Hudson, Kreitman, Aguadé (HKA) test (Hudson et al., 1987, Ferretti et al., 2012). Examination of relative levels of sequence polymorphism to divergence of the non-coding and protein-coding regions of these 2,482 genes using the MK test identified 472 non-coding and 217 protein-coding regions with signatures of positive selection. These non-coding regions include 18 known genetic determinants of *A. fumigatus* virulence, such as *zrfB* (plasma membrane zinc transporter), *myoE* (class V myosin, involved in cellular morphogenesis), and *pld2* (putative phospholipase D protein). The HKA test identified 207 non-coding and 4 protein-coding regions whose polymorphism to divergence ratio differed from a neutral locus. The 207 upstream non-coding regions identified with the HKA test included 4 genetic determinants of *A. fumigatus* virulence; 1 of the 4 protein-coding regions was a known genetic determinant of virulence. Molecular function terms enriched for genes associated with the non-coding regions that showed evidence of selection in both the MK and HKA tests, include ion-binding, transcriptional regulation, and stress response. These results demonstrate that *A. fumigatus* non-coding regions are typically more variable and more often under positive selection than their protein-coding counterparts, raising the hypothesis that they, too, may contribute to phenotypic differences between *A. fumigatus* strains.

## Methods

### Genomic data collection

All *Aspergillus* genomes are publicly available and were downloaded from NCBI (https://www.ncbi.nlm.nih.gov/); detailed information is provided in Table S1.

### Identification of single-copy orthologous genes

To infer single-copy orthologous genes across all 265 taxa, we used OrthoFinder, version 2.4.0 (Emms & Kelly, 2015). OrthoFinder clustered genes into orthogroups from sequence similarity information obtained using the program DIAMOND version 2.0.9 (Buchfink et al., 2015) with the proteomes of the 263 *A. fumigatus* strains and 2 *A. fischeri* strains as input. Fungal proteomes were obtained from a previously published study (Steenwyk et al., 2022). Key parameters used during sequence similarity search include an e-value threshold of 1 x 10^-3^ with a percent identity cutoff of 30% and a percent match cutoff of 70%. We considered genes to be single-copy orthologs if they were within the cutoff thresholds and were present in all 265 taxa.

### Retrieval of non-coding regions

To identify highly conserved non-coding regions, we first retrieved the non-coding sequences directly upstream of the first codon of all single-copy orthologous genes from all genomes. Non-coding sequence retrieval was performed using a custom Python script, which can be found at https://github.com/alecbrown24/General_Bio_Scripts (adapted from https://github.com/shenwei356/bio_scripts). We retrieved the first 1,500 base pairs (bp) of non-coding sequence directly upstream of the first codon of each gene and used these sequences to generate FASTA files of non-coding regions, as well as FASTA files of single-copy orthologous protein-coding sequences using Python version 3.8.2. For some non-coding regions, there were <1,500 bp of non-coding sequence between the first codon of the gene of interest and an upstream gene; in these instances, only the intergenic region was used for subsequent analyses.

### Alignment and identification of conserved non-coding and protein-coding regions

Multiple sequence alignments for all non-coding and protein-coding regions were constructed using MAFFT, version 7.453, with default parameter settings (Katoh et al., 2002). Codon-based alignments were inferred from the corresponding protein sequence alignments using pal2nal, version 14 (Suyama et al., 2006). Sequence identity in protein-coding and non-coding regions was calculated from their corresponding multiple sequence alignment files using AliStat version 1.12 (Wong et al., 2020). The percent sequence identity for each position in the alignment was calculated from the fraction of sites with the same nucleotide across all taxa. Sequence identity for protein-coding and non-coding regions can be found in Tables S2 and S3, respectively.

To measure evolutionary conservation across individual alignment sites, we implemented the PhyloP program as part of the Phylogenetic Analysis with Space/Time Models (PHAST) suite of programs (Ramani et al., 2019). PhyloP scores reflect the evolutionary conservation of individual nucleotide sites relative to the degree of conservation expected under neutrality. A positive score is predictive of evolutionary conservation and a negative score is predictive of evolutionary acceleration relative to neutral expectations.

### Phylogenetic tree inference

A phylogenetic tree of the 265 strains used in this study was generated by pruning from a larger *Aspergillus* species phylogeny (Steenwyk et al., 2022) using the Treehouse software in R using default parameters (Steenwyk & Rokas, 2019). Individual protein-coding region and non-coding region trees were inferred using IQ-TREE, version 2.0.6 (Minh et al., 2020), with “GTR+I+G+F” as it was the best fitting substitution model (Waddell and Steel, 1997; Vinet and Zhedanov, 2011).

### Identifying signatures of selection in *A. fumigatus* protein-coding and non-coding regions

To examine signatures of selection in *A. fumigatus* protein-coding and non-coding region alignments, we examined variation in the ratio of non-synonymous / non-coding sites (likely under selection) to synonymous sites (likely neutral) (**Fig. 1**). We used the protein-coding and / or non-coding region alignments to calculate the fractions of polymorphic (differences between *A. fumigatus* strains) and divergent sites (differences between *A. fumigatus* and the outgroup *A. fischeri*) for non-synonymous, synonymous, and non-coding sites using the standard McDonald Kreitman test function as part of the iMKT software in R (Murga-Moreno et al., 2019). For protein-coding regions, the ratio of polymorphic non-synonymous to synonymous sites was compared to the ratio of divergent non-synonymous to synonymous sites. For non-coding regions, the ratio of polymorphic non-coding to synonymous sites was compared to the ratio of divergent non-coding to synonymous sites (**Fig. 1**). For each MK test, the null hypothesis (H0) assumed that the ratio of selected vs neutral divergent sites was similar to the ratio of selected vs neutral polymorphic sites. We compared H0 to an alternative hypothesis (H1) in which there are more divergent sites than polymorphic sites across a given protein-coding or non-coding region, indicating positive selection. To determine whether H1 was significantly different from H0 for each of the codon-based alignments, we used Fischer’s exact test with a statistical significance threshold of p < 0.05 and a Bonferroni-adjusted alpha value < 0.01 to adjust for multiple testing. Results for both protein-coding and non-coding regions can be found in Table S4.

**Figure 1.**
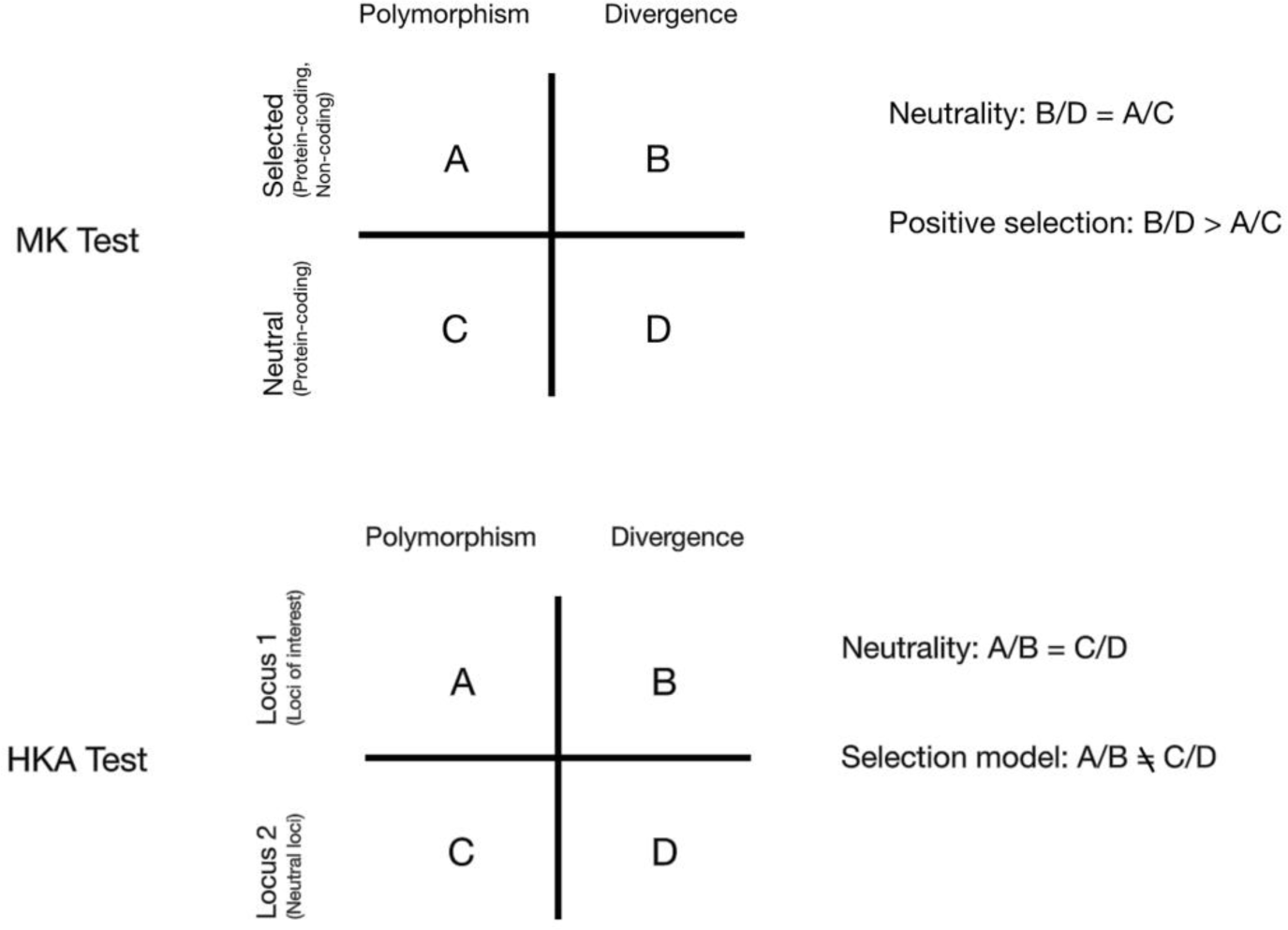
Brief overview of the MK and HKA tests of selection. The MK test (top) compares the polymorphisms (i.e., sites that vary within *A. fumigatus*) and divergence (sites that are fixed within *A. fumigatus* but differ from *A. fischeri*) between selected sites (non-synonymous or non-coding) and neutral sites (synonymous) between the protein-coding and non-coding regions of a given gene. For protein-coding regions, non-synonymous sites are compared to synonymous sites, while for non-coding regions, all sites are considered non-synonymous sites and are compared to the synonymous sites of the associated protein-coding region. Under a neutral model, the ratio of selected and neutral sites that are polymorphic is the same as the ratio of selected and neutral sites that are divergent. When the ratio of divergence is greater than the ratio of polymorphism, the MK test assumes that the selection is acting to fix advantageous non-synonymous changes, resulting in positive selection. The HKA test (bottom) compares the levels of polymorphism and divergence between two loci (the locus of interest and a reference, neutral locus). When ratio of polymorphism within species is equal to the ratio of divergence between species in the two loci, both loci are evolving neutrally. Should these ratios differ, we conclude that selection is occurring at the locus of interest.

The HKA test was also implemented, which compares the rate of polymorphism within *A. fumigatus* to divergence (between *A. fumigatus* and *A. fischeri*) at multiple loci (Hudson et al., 1987) (Fig. 1). The HKA test assumes that if two loci are evolving neutrally, the ratio of polymorphism to divergence at these loci should be relatively constant. We compared loci of interest to neutral loci using the HKADirect program (Ferretti et al., 2012). Neutral loci were determined by comparing each of the 2,482 loci to the genomic background using Tajima D’s test as part of the HKADirect program. The null hypothesis (H0) assumes that the patterns of genetic variation within a species (polymorphism) and the patterns of genetic differentiation between species (divergence) are consistent with neutral evolution. Under these conditions, the polymorphism to divergence ratio is similar between the loci of interest and neutral loci. We compared H0 to an alternative hypothesis (H1) in which assumes that the patterns of polymorphism and divergence at the loci of interest deviate from that of neutral loci due to the action of natural selection (Fig. 1). We used Fischer’s exact test with a statistical significance threshold of p < 0.05 to determine significance. Results for both protein-coding and non-coding regions can be found in Table S5.

Unlike the MK test, whose results can be used to directly compare the protein-coding and the non-coding regions of each gene, the HKA test instead compares each protein-coding and non-coding region to neutral loci. Thus, the MK test was used to determine differences in signatures of selection between non-coding regions and their associated protein-coding regions while the HKA test was used to detect signatures of selection in specific non-coding / protein-coding regions when compared to neutrally evolving non-coding / protein-coding regions.

### Functional enrichment analyses of genes with signatures of selection

To determine whether genes with signatures of selection in either their protein-coding or non-coding regions were enriched for particular functional categories, we implemented the Gene Ontology (GO) tool g:PROFILER (Raudvere et al., 2019) using default settings. We performed four separate analyses of functional enrichment among genes that significantly differed from the null hypothesis based using the MK test of protein-coding and non-coding regions as well as the HKA test of protein-coding and non-coding regions. Each of these gene sets was compared to a general background set that includes all the features / gene names in the Ensembl genomes database with at least one GO annotation for *A. fumigatus*. All functional enrichment analyses used a p-value cutoff of 0.05. All genes found to be statistically significant can be found in Table S6.

### Examination and visualization of mutational signatures

To identify interesting examples of sequence variation between *A. fumigatus* strains for non-coding regions of genes of interest, we visualized and compared multiple sequence alignments using the MView function in EMBL-EBI (Madeira et al., 2019). Workflow of methods can be seen in Supplemental Figure 1.

## Results

### Protein-coding and non-coding regions exhibit differing levels of sequence conservation within *A. fumigatus*

To analyze the sequence diversity of non-coding regions across *A. fumigatus* strains (Fig. 2), we first identified 5,812 single-copy orthologous genes amongst 263 *A. fumigatus* strains and 2 *A. fischer*i strains. We then measured evolutionary conservation at individual alignment sites for protein-coding and non-coding regions across the 5,812 single-copy orthologous genes of interest. Of the 5,812 single-copy orthologous genes, 5,646 were found to be alignable; thus, we focused our subsequent analyses around these genes.

**Figure 2.**
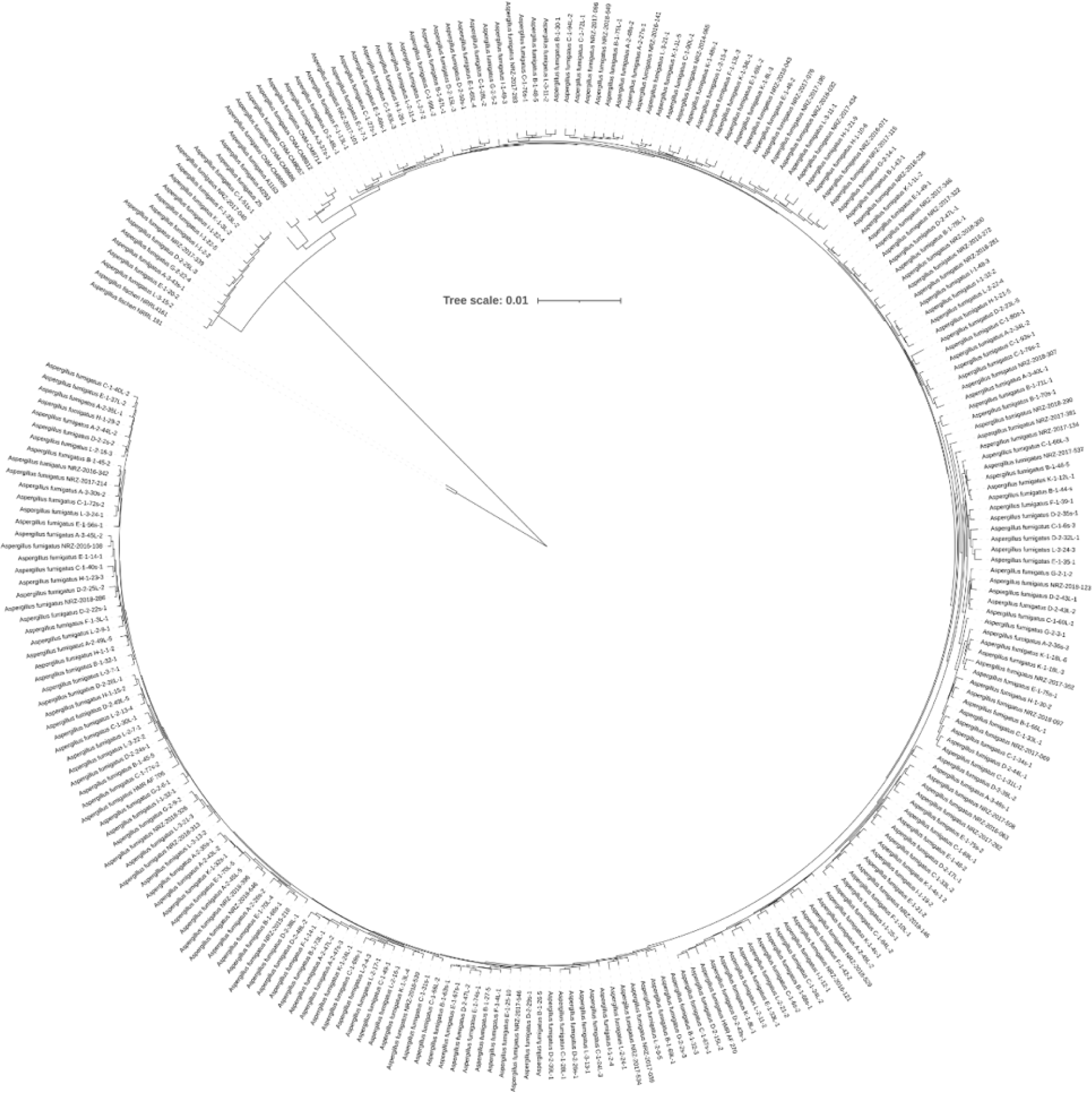
Phylogeny of the 263 A. *fumigatus* strains used in this study. A phylogenetic tree of the 265 strains used in this study was generated by pruning from a larger *Aspergillus* species tree inferred from analyses of 1,362 protein-coding regions (Steenwyk et al., 2022).

Examination of PhyloP scores for each protein-coding and non-coding alignment individually revealed that both protein-coding and non-coding regions exhibit varying levels of conservation across single-copy orthologous genes (Fig. 3). Examination of average PhyloP scores in protein-coding regions revealed a lower area of conservation near their start (first ∼100 bp). This may be due to slight differences in gene annotation between strains or the presence of genuine variation; utilization of alternate start sites for the same gene has been demonstrated in *Aspergillus* (Kjærbølling et al., 2020). Beyond the first ∼100 bp, conservation levels of protein-coding regions remain high throughout the first 1,500 bp. High conservation among protein-coding regions is also consistent with comparisons between *A. fumigatus* and closely related species (Fedorova et al., 2008).

**Figure 3.**
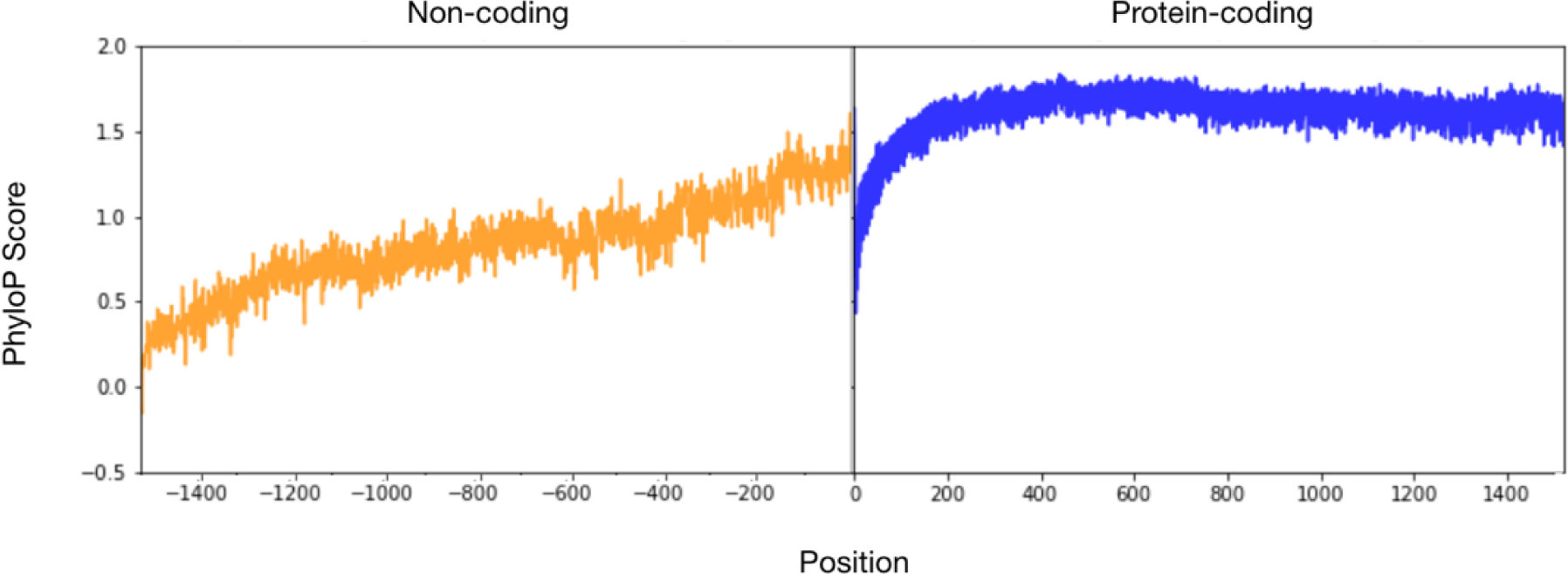
Non-coding regions are less conserved than protein-coding regions in the major fungal pathogen *A. fumigatus*. PhyloP Score for protein-coding and non-coding regions. To determine conservation in protein-coding (blue) and non-coding (orange) regions, we calculated the PhyloP score across sites of all multiple nucleotide sequence alignments of protein-coding and upstream non-coding regions of 5,646 single copy orthologs across 263 A*. fumigatus* strains and 2 A*. fischeri* strains. Scores of conservation were measured for individual nucleotide sites and scores were then averaged across all orthologs. Conserved sites have PhyloP scores above 0 and non-conserved sites have scores below 0. We find that in non-coding regions, sites that are closer to the start of the transcription start site (TSS) exhibit a higher level of conservation and generally decrease in conservation as we move further from the TSS. In protein-coding regions, PhyloP scores are generally above 0, which are indicative of high sequence conservation; the lowest scores are observed near the start of the protein-coding regions, which is likely an artifact caused by variation in starting codon position of gene annotations across *A. fumigatus* strains.

The average PhyloP score in non-coding regions across *A. fumigatus* strains revealed that the highest levels of sequence conservation (as indicated by a higher PhyloP score) were directly upstream of the start site, with conservation generally decreasing further away from the start site. We also found that conservation begins to fade around 1,500 bp upstream of the start site (as indicated by a PhyloP score of 0). This pattern of higher sequence conservation in non-coding regions right upstream of the transcription start site is consistent with a previous study of non-coding regions comparing A. *fumigatus* and closely related species (Brown et al., 2022).

To further examine the conservation of pairs of non-coding and protein-coding sequences, we calculated the percent identity for all single-copy orthologs in their protein-coding and associated non-coding regions. We found that percent nucleotide sequence identity of protein-coding regions exhibited weak but significant correlation (r^2^ = 0.143 and p-value < 0.0001) with the percent nucleotide sequence identity of the associated non-coding regions (i.e., orthologs with higher percent identity in their protein-coding regions also exhibited higher percent identity in their non-coding regions). Thus, it was sometimes the case that highly similar protein-coding regions were associated with non-coding regions that display higher sequence variation. This result suggested that the functions of genes with highly conserved protein-coding regions may still differ between strains due to differences in the genes’ non-coding regions. In a few cases, we also identified highly divergent protein-coding regions that were associated with highly similar non-coding regions.

We next computed the percent nucleotide sequence identity between the non-coding and protein-coding regions of each single-copy orthologous gene across all 265 strains (Fig. 4) for the 5,646 genes whose protein-coding sequences were alignable. Averaging the non-coding region percent similarities for the 5,646 single-copy orthologous genes revealed an average identity of ∼85%, while the average protein-coding percent identity was ∼92%. Additionally, we found that 928 protein-coding alignments exhibited <75% nucleotide sequence identity (this number includes the 166 unalignable genes), 2,482 exhibited sequence identity between ≥75% and <99%, and 2,402 exhibited ≥99% identity. For non-coding region alignments, 1,274 non-coding alignments exhibited <75% identity, 3,721 that exhibited sequence identity between ≥75% and <99%, and 817 that exhibited ≥99% identity.

**Figure 4.**
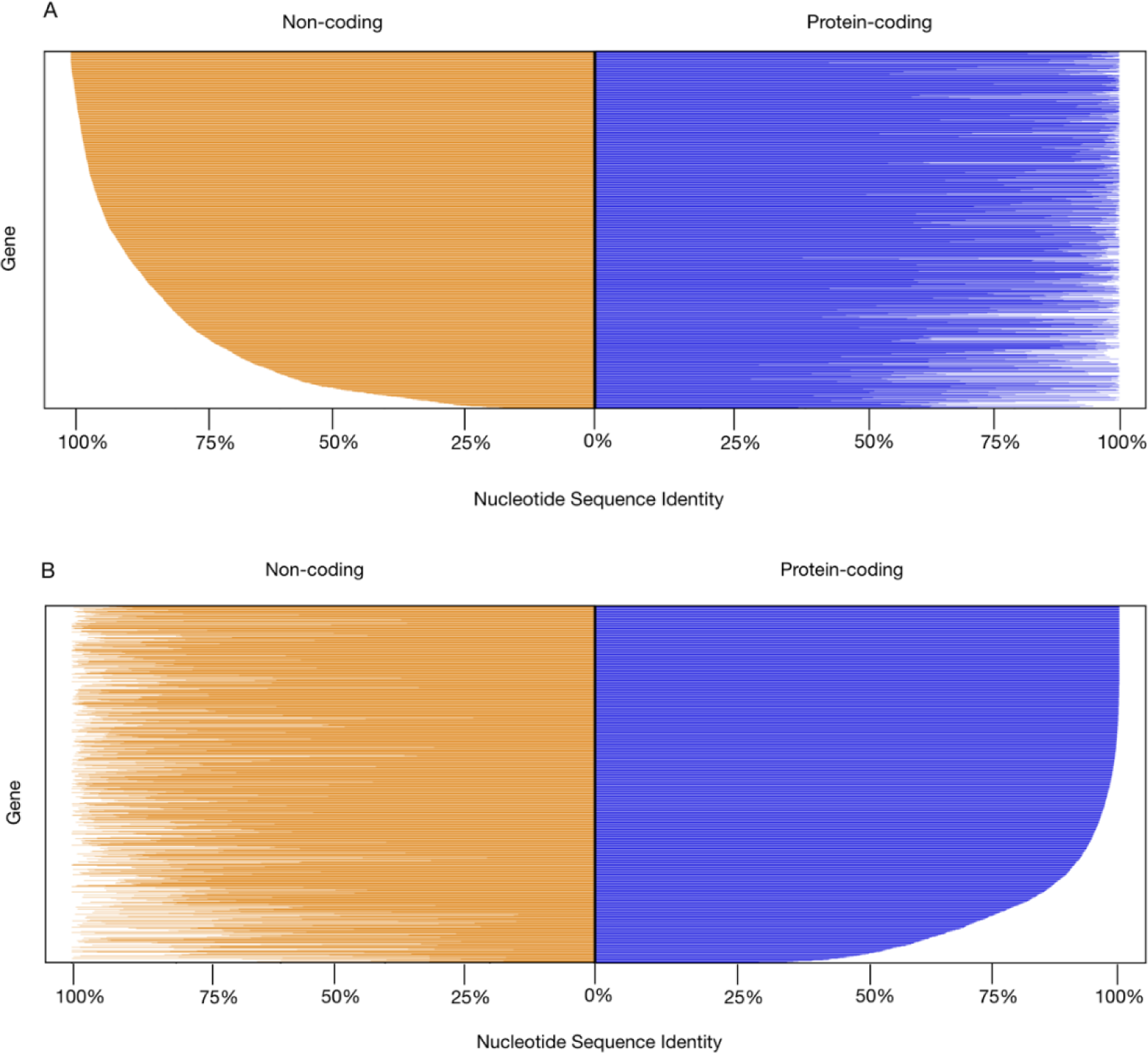
*A. fumigatus* orthologs exhibit numerous instances of highly conserved protein-coding genes whose non-coding regions are poorly conserved. Percent identity of protein-coding and non-coding regions of 5,646 *A. fumigatus* genes. Up to 1.5kb upstream noncoding region (orange) is shown to the left were calculated and plotted by percent identity starting with 100% similar (top row) and descending. The associated protein-coding regions (blue) are shown to the right. Although the sequence conservation of protein-coding regions and the sequence conservation of their corresponding non-coding regions are correlated, there are numerous instances of genes with high protein-coding sequence identity and a lower identity in their non-coding region. A) *A. fumigatus* genes ranked by percent nucleotide sequence identity of their non-coding regions. B) *A. fumigatus* genes ranked by percent nucleotide sequence identity of their protein-coding regions.

### Many non-coding regions have signatures of positive selection

To examine signatures of selection in *A. fumigatus* genes, we performed the MKA and HKA tests of selection in 2,482 pairs of protein-coding and non-coding region alignments. For the MK test, we found that a total of 472/2,482 (19.0%) genes exhibited signatures of selection in their non-coding regions but not in their protein-coding regions, a total of 217/2,482 (8.7%) genes experienced selection in their protein-coding regions but not in their non-coding regions, and 144/2,482 (5.80%) genes experienced selection in both their protein-coding and non-coding regions (Fig. 5).

**Figure 5.**
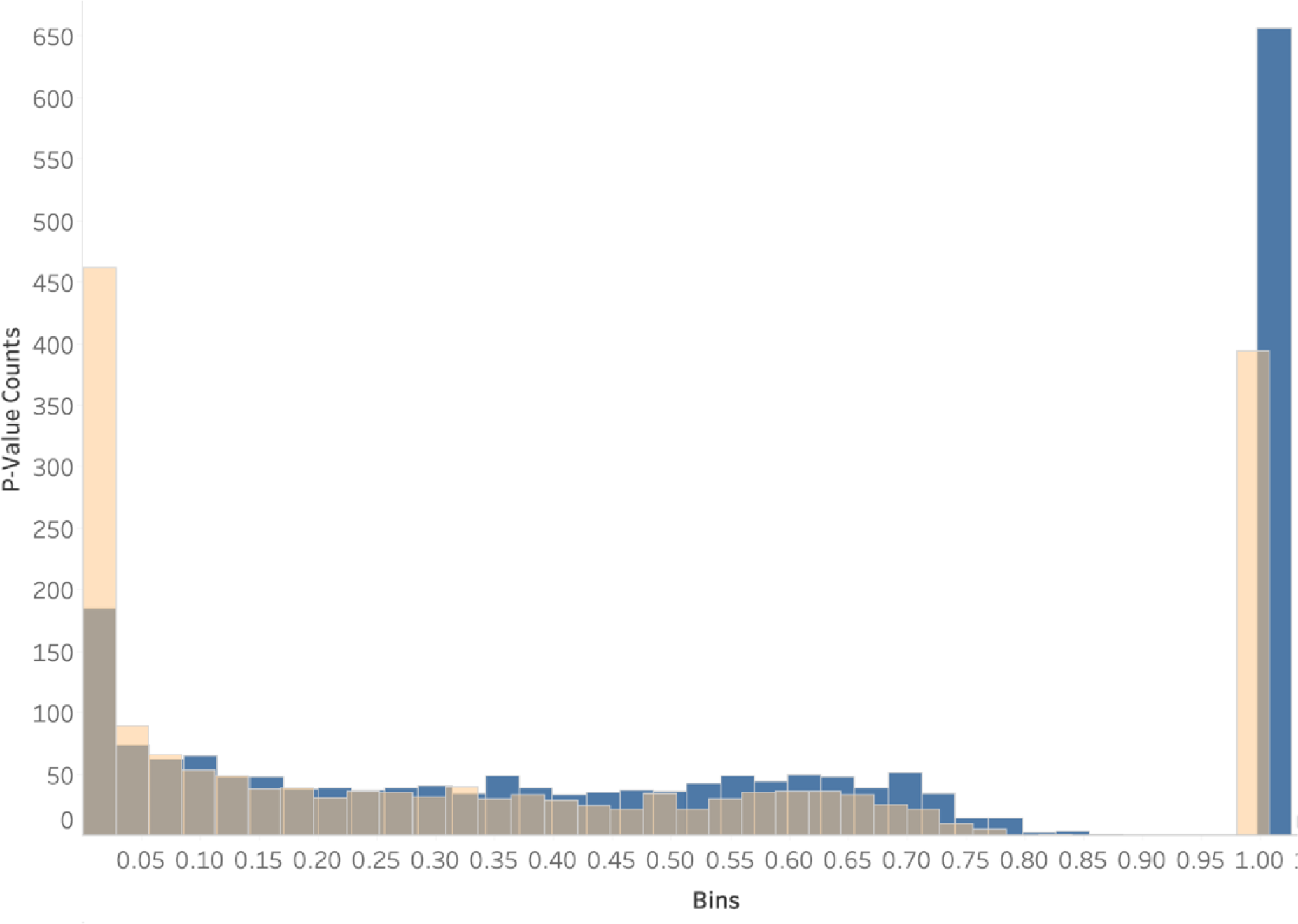
A higher number of non-coding regions than protein-coding regions exhibit signatures of selection under the McDonald-Kreitman test. Histogram of the distribution of p-values of the MK test. The MK test was calculated for 2,482 single copy orthologs in both protein-coding (blue) and non-coding regions (orange). 217 protein-coding and 472 non-coding regions were found to be significant (p < 0.05).

For the HKA test, we found 4/2,482 (8.8%) and 207/2,482 genes with evidence of positive selection in their protein-coding and non-coding regions, respectively (Fig. 6). Examination of the genes that were significant under the MK and HKA tests shows that there relatively limited overlap for both protein-coding and non-coding regions (Fig. 7). For example, only 36 non-coding regions exclusively exhibit evidence of selection by both tests. For protein-coding regions, the lack of overlap is largely due to the very small number of protein-coding regions that show evidence of selection in the HKA test. For the non-coding regions, the limited overlap is likely due to the differences in the neutral sites used by the two tests (the MK test uses the synonymous sites of the corresponding protein-coding region whereas the HKA test uses all the sites of a neutrally evolving non-coding region).

**Figure 6.**
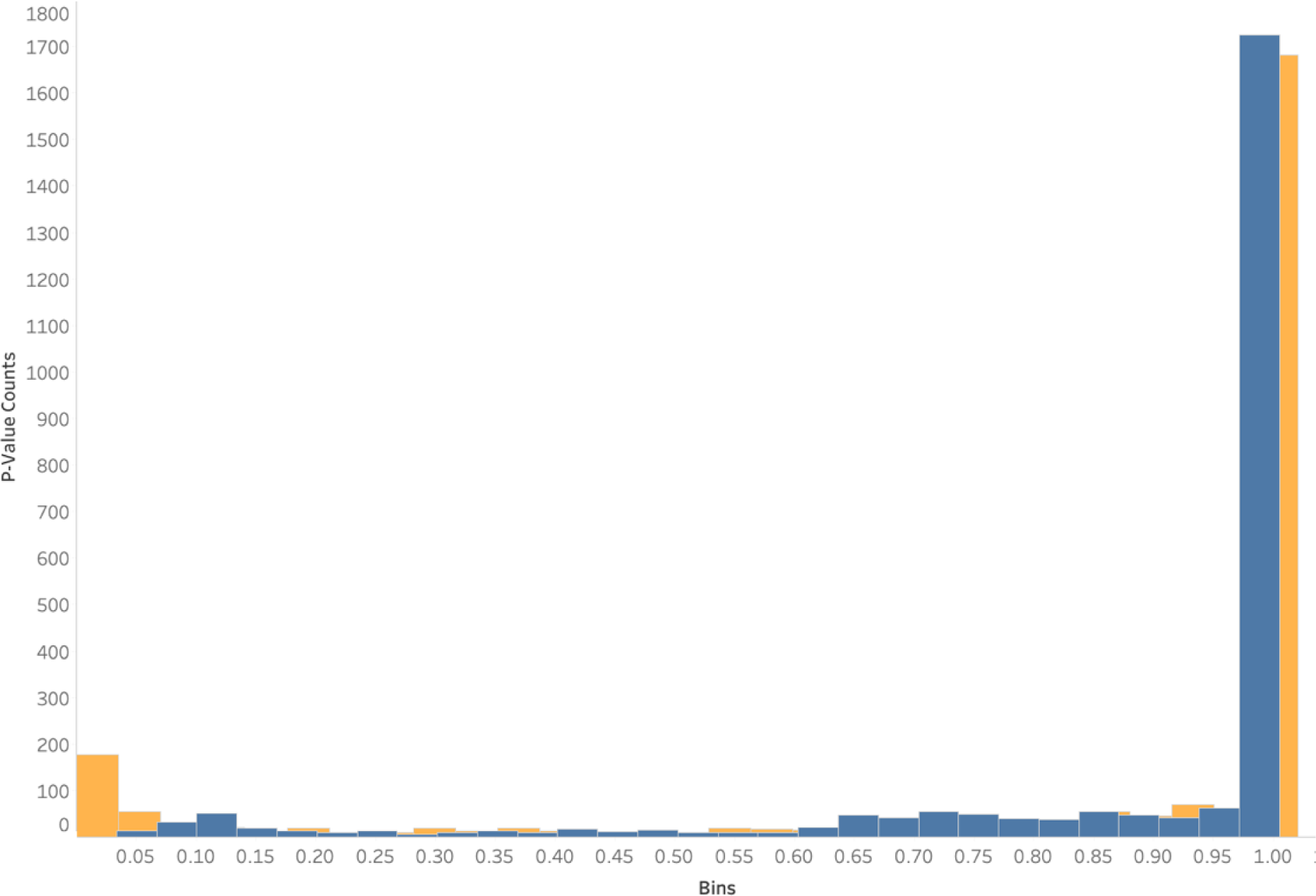
HKA test identified 207 non-coding and 4 protein-coding regions that exhibit a signature of selection. Histogram of the distribution of p-values of the HKA test across protein-coding (blue) and non-coding (orange) regions of 2,482 genes. 207 non-coding and 4 protein-coding regions were found to be significant (p < 0.05).

**Figure 7.**
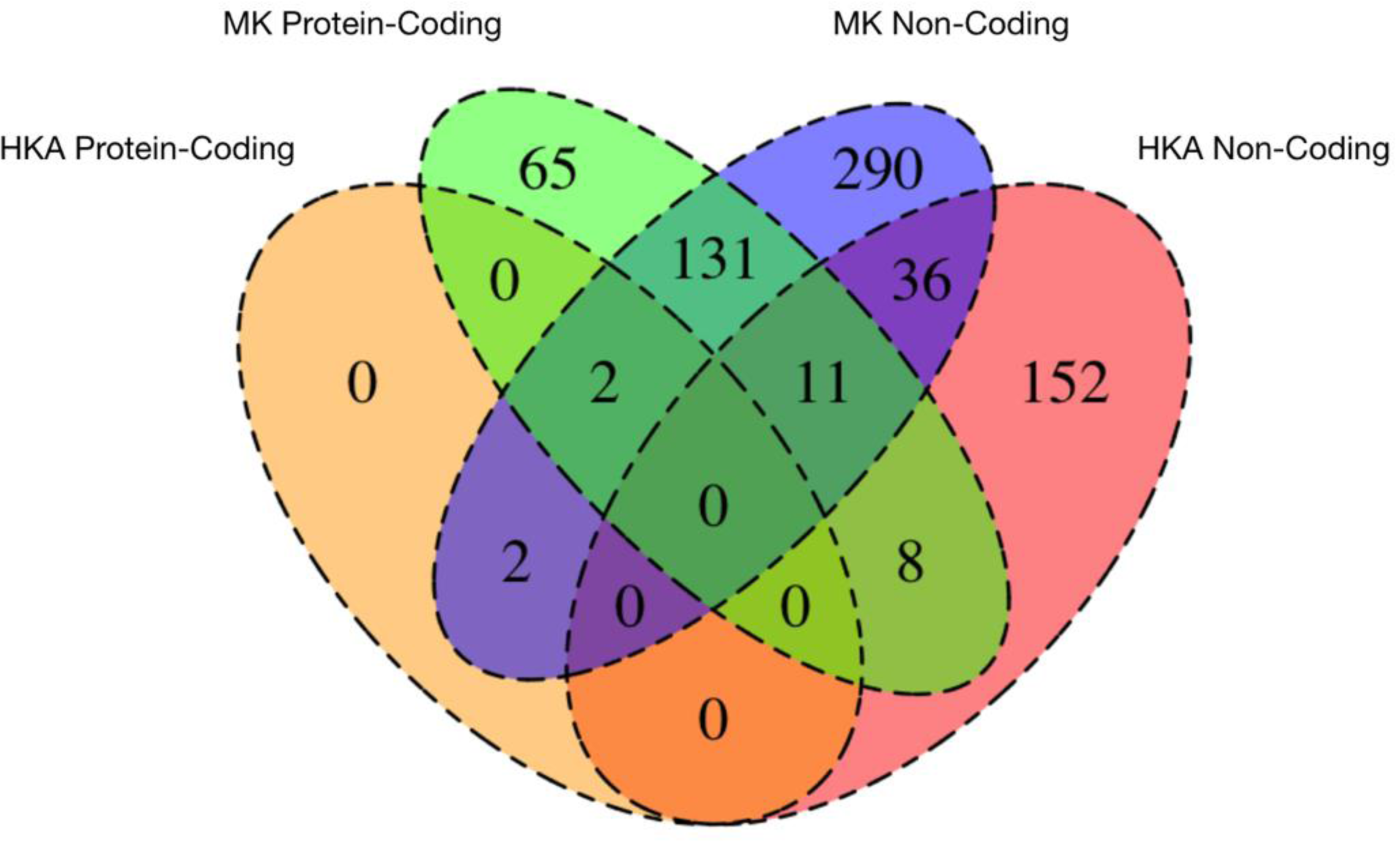
Venn Diagram of significant results from HKA protein-coding, MK-protein-coding, MK non-coding and HKA protein-coding tests. There were 478 (290 + 36 + 152) genes with evidence of selection only in their non-coding regions compared to 65 genes with evidence of selection only in their protein-coding regions across both MK and HKA tests.

### Genes with evidence of selection in non-coding regions are enriched for binding and regulatory activity, including 21 genes involved in *A. fumigatus* virulence

We used GO enrichment to determine if any functions were overrepresented. For the 472 genes with evidence of selection under the MK test in their non-coding regions, we found 22 categories that were enriched for molecular function. “DNA Binding” was the top term identified for molecular function (p=9.06×10^-8^) and the most enriched term overall. “Cytoskeleton motor activity (p=6.59×10^-6^), “Ion Binding” (p=7.33×10^-5^) and “Transcription factor binding” (p=4.08×10^-4^) were also represented grouped terms for molecular function components respectively (Table S7). For the 217 genes with evidence of selection in their protein-coding regions, 7 molecular functions were overrepresented, including “ATP hydrolysis activity” (p=1.37×10^-3^) and various functions involved in binding activities. For the HKA test, we find that enzyme regulator activity (p =3.33×10^-2^) was the only molecular term found for the non-coding regions, and no GO terms were enriched for the HKA protein-coding results (Table S7). Additionally, the 36 genes that experienced selection in their non-coding regions under both the HKA and MK tests, are enriched for various regulatory processes (Table S7).

We compared our list of genes under selection to a previously curated set of 206 genetic determinants of *A. fumigatus* virulence (Steenwyk et al., 2021). Given that *A. fumigatus* strains have been demonstrated to exhibit differences in virulence in mouse models of fungal disease (Keizer et al.,2021), selection in the non-coding or protein-coding regions of these genes may be of relevant to *A. fumigatus* virulence. We found that the non-coding regions of 18 of the 206 virulence genes were under selection according to the MK test (*argEF, ags1, csmB, pabA, medA,mtfA ,myoB ,myoE ,pld2,* g*liP, rgsC, aceA, atfA, cch1, fbx15, flcB, schA, zrfB)* and 3 according to the HKA test *(cds1*, *nop4*, *dvrA*). We found that the most represented general function amongst these 18 genes was “stress response”, which raises the question of their impact on virulence, given the role that stress response has been shown to play in *A. fumigatus* virulence (Colabardini et al., 2022); for example, exposure to different temperatures, pH and drug treatments lead to differences in gene expression (Latge et al., 2019; Colabardini et al., 2022). We also identified 4 protein-coding regions of virulence genes (*lysF*, *myoB*, *aftA*, *fbx15*) that showed evidence of selection under the MK test. Similarly, we found 11 virulence genes (*hcsA*, *lysF*, *ags1*, *chsG*, *erg12*, *rtfA*, *tom24*, *fmaE*, *gliC*, *gliT*, *sidI*) with evidence of selection in their non-coding regions under the HKA test, and 1 (*aceA*) with evidence of selection in its protein-coding region.

### Examples of non-coding region differences between *A. fumigatus* strains

We next sought to identify representative sequence differences in non-coding regions between *A. fumigatus* strains that exhibited signatures of selection according to the MK test or HKA test. (Fig. 8). One such example was the non-coding region of *AFUA_3G03310*, which is under selection according to the MK test. The non-coding region exhibits a 12 bp region (AGCCACAGAACT) present in *A. fumigatus* Af293 and 91 other strains but absent from the rest. This binding site location is an exact match to the Met31 transcription factor binding site involved in sulfur metabolism in *S. cerevisiae* (Cormier et al., 2010). The MetZ transcription factor performs a similar function in *Aspergillus nidulans* (Pilsyk et al., 2015) and may be also in *A. fumigatus* (Amich et al., 2016). Another gene with evidence of selection in its non-coding region is *AFUA_3G11330*, which encodes the putative transcription factor AftA involved in stress response and spore viability in *Aspergillus* (Lara-Rojas et al., 2011). The *AFUA_3G11330* non-coding region exhibits a 6 bp region (TACTCT) present in *A. fumigatus* Af293 and about half of the other *A. fumigatus* strains while absent in the other *A. fumigatus* strains. This 6 bp region is similar to the 6 bp binding site for Yap1 in *S. cerevisiae*, which is required for oxidative stress tolerance (Natkanska et al., 2017). An ortholog of Yap1 is known to be involved in voriconazole resistance in *A. flavus* (Ukai et al., 2018) and may play a role in stress response in *A. fumigatus*, which is important as a mechanism for survival within the human lung (Latge et al., 2019). Both the MK and HKA tests found signatures of selection in *AFUA_6G03660*, an uncharacterized gene in *A. fumigatus*. Although of unknown function, this gene is of particular interest as its non-coding region differs between the two reference strains *A. fumigatus* Af293 and *A. fumigatus* A1163, which vary in their virulence in animal models of fungal disease. *A. fumigatus* A1163 exhibits a larger region of 22 bp that is absent in *A. fumigatus* A1163. This region helps to illustrate the complex non-coding sequence differences between *A. fumigatus* strains, including those that are closely related.

**Figure 8.**
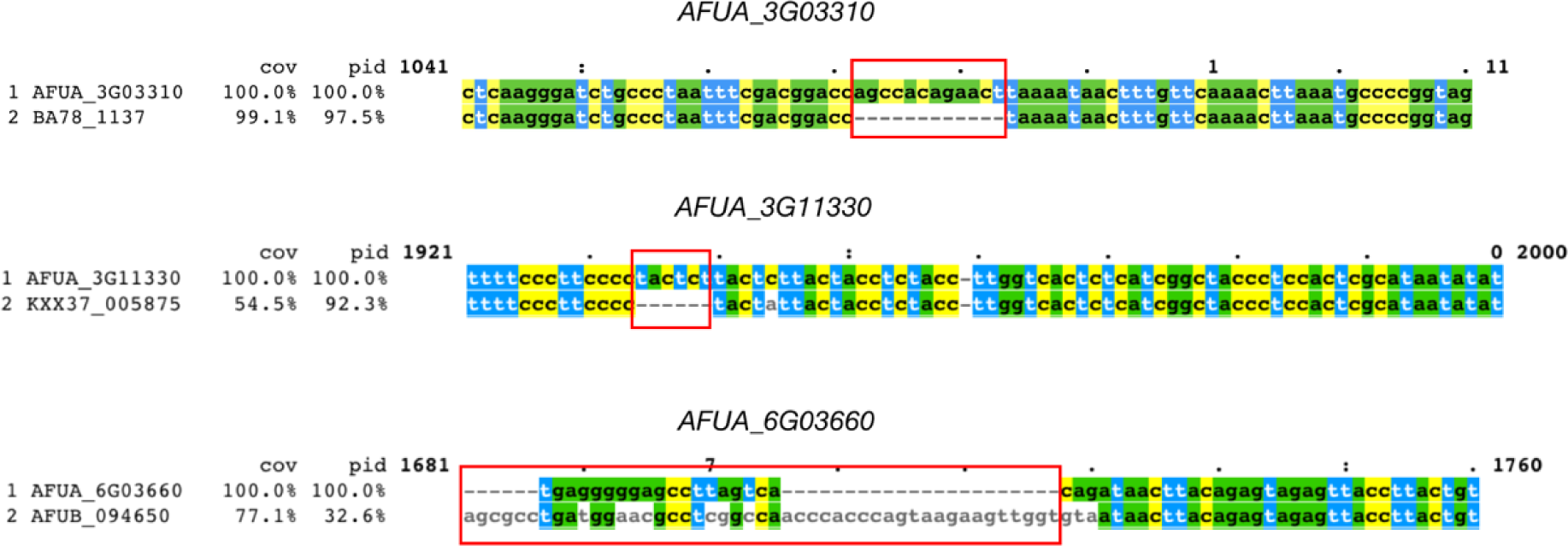
Notable examples of sequence differences between A. *fumigatus* strains in non-coding regions. Here, we show three regions of sequence alignments of non-coding regions that differ between *A. fumigatus* Af293 and another *A. fumigatus* strain. (A) AFUA_3G03310 (RTA1 domain protein) exhibits a 12 bp region present in *A. fumigatus* Af293 but absent in several other strains, including BA78_1337. (B) AFUA_3G11330 (transcription factor AtfA) exhibits a 6 bp region absent in *A. fumigatus* Af293 but present in several other strains, including KXX37_005875. (C) AFUA_6G03660 (predicted to be involved in the production of biotin) exhibits a non-coding difference between the two reference strains of *A. fumigatus* (Af293 and A1163).

## Conclusion

Here, we presented a comprehensive study of signatures of positive selection in both protein-coding and non-coding regions across many strains from the fungal pathogen *A. fumigatus*. We identified several non-coding regions under selection in *A. fumigatus*, including several candidate transcription factor binding sites that differ between strains are await further exploration. Currently, there are no datasets available that report genome-wide differential expression data for *A. fumigatus* strains. Experiments on diverse *A. fumigatus* strains to investigate and analyze the differential expression of genes and examine the functional implications of the non-coding region disparities we have identified, specifically in relation to the expression and virulence of *A. fumigatus*, will be of great interest. More broadly, these findings suggest divergence in non-coding sequences may play an important role in population variation among fungal pathogens and beyond.

## Data Availability

All *Aspergillus* genomes are publicly available and were downloaded from NCBI (https://www.ncbi.nlm.nih.gov/).

## Acknowledgements

We thank members of the Rokas lab for helpful discussions and feedback. Research in A.R.’s lab is supported by grants from the National Institutes of Health National Institute of Allergy and Infectious Diseases (R01 AI153356), the National Science Foundation (DEB-2110404), and the Burroughs Wellcome Fund. The content is solely the responsibility of the authors and does not necessarily represent the official views of the National Institutes of Health. JLS is a Howard Hughes Medical Institute Awardee of the Life Sciences Research Foundation.

